# Estradiol treatment in a nonhuman primate model of menopause preserves affective reactivity

**DOI:** 10.1101/248591

**Authors:** Eliza Bliss-Moreau, Mark G. Baxter

## Abstract

As humans age, their affective lives tend to become more positive and less negative. This phenomenon, known as the positivity effect (or positivity bias), occurs even as aging leads to declines in health and cognitive outcomes. Despite these well documented effects in humans, extent to which affective processes change in nonhuman animals, and in particular nonhuman primates – is unclear. As a first step towards developing a model for human affective aging in rhesus monkeys (*Macaca mulatta*), we tested aged, surgically menopausal aged and middle-aged gonadally intact female rhesus monkeys on a classic index of affective reactivity in monkeys, the Human Intruder task. The Human Intruder task evaluates behavioral responses to varying levels of threat. Aged, surgically menopausal monkeys received hormone replacement therapy consisting of a cyclic estradiol regimen, or vehicle injections as a control. Average responsivity to threat did not vary by condition, but middle aged monkeys and aged monkeys on estradiol were more reactive to the most potent level of threat than to a moderate level of threat, replicating previously published results in other age groups and male monkeys. In contrast, aged monkeys not on estradiol did not show such calibration to threat level. These findings suggest that estrogen may be important for maintaining more youthful affective responding. They also illustrate the utility of behavioral assays of affective reactivity in nonhuman primate models of cognitive and reproductive aging in humans.

## Introduction

Something curious happens to affective life as people age – it gets better. On the whole, affective life becomes more positive and less negative as people traverse the time between middle age and old age, a pattern known as the “positivity effect” (for reviews Carstensen & Mikels, 2005; Mather & Carstensen, 2005). This change happens even in the context of the increased incidence of disease in older people (Chatterji, Byles, Cutler, Seeman, & Verdes, 2015), deleterious effects of aging on memory (Raz, 2000), and the shrinking of social networks with advanced age (Cornwell, Laumann, & Schumm, 2008; McPherson, Smith-Lovin, & Brashears, 2006). Variation in affective life across older adulthood is evident both in instrumental tasks (e.g., those that measure attention) and also self-reports of emotional experience. For example, accumulating evidence demonstrates that the visual attention of older adults, unlike younger adults, is biased towards positive information and, or, away from negative information (e.g., Frewen, Dozois, Joanisse, & Neufeld, 2008; Isaacowitz, Wadlinger, Goren, & Wilson, 2006a, 2006b; Mather & Carstensen, 2003; Nikitin & Freund, 2011). Older people’s emotional well-being is greater than younger people’s, either as a result of a reduction in negative emotional experiences and, or, an increase in positive emotional experiences. For example, during a 1-week experience sampling study, there were no age differences in the frequency of positive emotional experiences or the intensity of either positive or negative experiences, but older adults reported significantly fewer negative emotional experiences than did middle-aged adults (Carstensen, Pasupathi, Mayr, & Nesselroade, 2000). Follow-up study of those participants 5 and 10 years past the original assessment indicated that positive emotional experiences increased into the late 60s, after which no changes were observed (Carstensen et al., 2011).

Despite the robustness of the positivity effect in humans, the biological mechanisms that support it are largely unknown. One possibility is that the positivity effect is subserved, at least in part, by changes in hormone production, especially for women who experience menopause. Estrogen has been widely implicated in affective processes (for reviews Gasbarri, Tavares, Rodrigues, Tomaz, & Pompili, 2012; Georgakis et al., 2016; Newhouse & Albert, 2015; Sherwin, 1996), opening the possibility that significant changes to estrogen production in post-versus pre-menopausal women might play a role in observed changes in affective life. Estrogen levels appear to influence the perception of some facial displays of emotion (Guapo et al., 2009; Kamboj, Krol, & Curran, 2015). Estrogen levels impact mood (for a review Newhouse & Albert, 2015). For example, following hysterectomy and ovariectomy, women receiving monthly estrogen (specifically, estradiol) injections reported less negative affect and more positive affect when estradiol levels were higher (Sherwin & Disord, 1988). Further, early menopause dramatically increases women’s likelihood of clinical depression (Georgakis et al., 2016).

One approach to understanding the mechanisms that generate psychological and behavioral processes – such as the positivity effect – is to adopt an animal model that affords precise experimental control and the ability to manipulate biology to map causation. Nonhuman primates, and in particular, rhesus monkeys (*Macaca mulatta*), the most widely used nonhuman primates in research (Carlsson, Schapiro, Farah, & Hau, 2004), represent a tremendous resource for such study given their complex social and affective behavior and neuroanatomy. As a first step in the development of a nonhuman primate model of normal, healthy affective aging, we carried out a study investigating affective reactivity to threat in a group of aged rhesus monkeys who had undergone surgical menopause and were either treated with hormone replacement therapy or not, and a group of middle aged female monkeys. Effects of estradiol on affect have been described in young monkeys in the context of both normal menstrual cycling (Lacreuse, Martin-Malivel, Lange, & Herndon, 2007), as well with estradiol replacement in ovariectomized young female monkeys (Lacreuse et al., 2007), but to our knowledge the impact of estradiol on affect has not been evaluated in nonhuman primates in the context aging.

Our goal in this initial study was to extend our well-characterized monkey model of cognitive aging and hormone replacement therapy (Baxter et al., 2013a; Hara et al., 2014; Morrison & Baxter, 2012; Rapp, Morrison, & Roberts, 2003) to the domain of affect. As a first step, we employed a well-validated affective processing task used widely with rhesus monkeys, the classic Human Intruder task, to evaluate affective reactivity to different levels of threat (Bliss-Moreau & Moadab, 2016; Gottlieb & Capitanio, 2013; Kalin & Shelton, 1989, 1998). We reasoned that if rhesus monkeys also experience the positivity effect, then aged menopausal animals should show reduced affective reactivity to threatening stimuli.

## Method

All experimental procedures carried out at the California National Primate Research Center (CNPRC). All protocols were approved by the University of California, Davis Institutional Animal Care and Use Committee.

### Subjects

Subjects were 24 adult female rhesus macaques, from two different experimental cohorts. Cohort 1 included aged animals from a study on the impact of hormone replacement therapy on cognition following surgically induced menopause N=17 (*M*_*age*_=22.02, *SD*_*age*_=1.18). Monkeys in Cohort 1 were ovariectomized approximately 2 years prior to data collection and, at the time of testing were either receiving hormone replacement therapy (N=6) or not (i.e., controls N=11), as described below. Cohort 2 included middle-aged, gonadally intact females N=7 (*M*_*age*_=12.65, *SD*_*age*_=3.08) who are part of a larger studying on affective reactivity.

Subjects in Cohort 1 were housed either singly (N=4, 1 in hormone treatment group) or with a social partner (N=13, 5 in hormone treatment group) in standardized primate caging. Animals that were socially housed had access to their social partner (and her cage) 8-hrs per day. Subjects in Cohort 2 were all pair-housed with a compatible vacestomized male partner either 24-hrs per day or 8-hrs per day in standard primate caging. In all cases, indoor housing rooms were maintained on a 12-hour light/dark cycle (lights on at 6 am). Animals were fed monkey chow (Lab Diet #5047, PMI Nutrition International INC, Brentwood, MO) twice daily, provided with fresh fruit and vegetables twice per week, had access to water ad libitum, and provided with a variety of enrichment activities regularly.

### Cohort 1: Ovariectomy Surgery and Hormone Treatment

Subjects in Cohort 1 underwent ovariectomies at an average of 20.18 years of age (*SD*=0.93). For ovariectomy (OVX) surgery, monkeys were sedated with 10 mg/kg ketamine i.m., given 0.04 mg/kg atropine s.c., then intubated, placed on isoflurane anesthesia to effect, and positioned in dorsal recumbency. A ventral caudal midline abdominal incision visualized the body of the uterus and both ovaries. Ovarian vessels and the fallopian tubes were isolated, ligated, and severed. The abdominal wall was then closed in three layers with two layers of 2/0 absorbable suture and the final subcuticular layer with 3/0 absorbable suture. The animals were recovered in the CNPRC surgical recovery unit and given three days of postoperative analgesia, 1.5 mg/kg oxymorphone i.m. three times daily. Daily urine assays for one month following OVX confirmed that animals were not spontaneously generating estrogen (for further details about hormone assays see Baxter et al., 2013b; Baxter, Santistevan, Bliss-Moreau, & Morrison, submitted).

Subjects in the hormone treated condition received injections of estradiol: 100 μg of estradiol cypionate in 1 ml peanut oil vehicle; 2 i.m. injections given 9 hours apart. Subjects in the control condition received injections of vehicle (peanut oil) at the same time intervals. Testing occurred 2 days after the previous hormone injection.

### Responsivity to Threat

Subjects completed a standard evaluation of behavioral responsivity to threat for nonhuman primates called the “Human Intruder” (HI) task (e.g., Bliss-Moreau & Moadab, 2016; Gottlieb & Capitanio, 2013; Kalin & Shelton, 1989, 1998). In the HI, subjects are placed in a standard primate cage (85.5 cm W x 68 cm D x 82 cm H, bottom of cage ~130 cm off of the ground) in a test room that houses no other monkeys. After a 1 minute acclimation phase, subjects are presented with an unfamiliar human (in this case a female experimenter, approximately 1.75 m tall) in each of four different positions (1 minute each): 1) *profile-far*: human standing 1 m from cage facing 90 degrees away from cage; 2) *profile-near*: human standing 0.3 m from cage facing cage facing 90 degrees away from cage; 3) *stare-far*: human standing 1 m from cage facing cage and making eye contact with subject; 4) *stare-near*: human standing 0.3 m from cage facing cage and making eye contact with subject. Behavior of the subjects was video-taped using a webcam (Logitech C920m) connected to a laptop computer to record the video located outside the testing room and monitored by an experimenter. Animals in Cohort 1 completed only 1 day of testing and animals in Cohort 2 completed 5 days of testing as part of a larger experiment. To equate samples, only data from the first day of testing in Cohort 2 were analyzed for this report.

### Behavioral Scoring

Behavior was scored using a standardized ethogram that recorded position in the cage and affect-related behaviors, using a 1/0 scoring method and 10 second bins. The scored affect-related behaviors were: lipsmack face, bared teeth face, threat face, freeze, coo vocalization, scream vocalization, grunt vocalization, bark vocalization, cage-shake (physical display), toothgrind, self-scratch, yawn, and stereotypy (including pace, spin, bounce, etc.). If a behavior was present during a given 10 second bin, that behavior received a score of 1 for that bin. If a behavior was not present for a given 10 second bin, that behavior received a score of 0 for that bin. Bins representing the generation of affect-related behaviors were summed into an index representing affective reactivity.

### Data Analysis Strategy

Data analyses were completed in SPSS 24.0. As described below, we computed the
bias corrected boot strapped mean and 95% confidence interval (CIs) for each metric of interest and compared the means and CIs to determine if there were within condition differences in HI reactivity.

## Results

Affective reactivity scores for both the aged and middle aged animals during the profile condition were essentially 0, regardless of the intruder’s distance from the cage. In contrast, all animals were reactive during the stare condition and the mean magnitude of that reactivity was essentially consistent across age and hormone conditions. Only aged hormone treated animals generated significantly greater affective reactivity to the near versus far condition based on the non-overlapping 95% confidence intervals. Variance in aged hormone treated animals was also less than in the other groups. See Figure 1, below.

**Figure 1:**
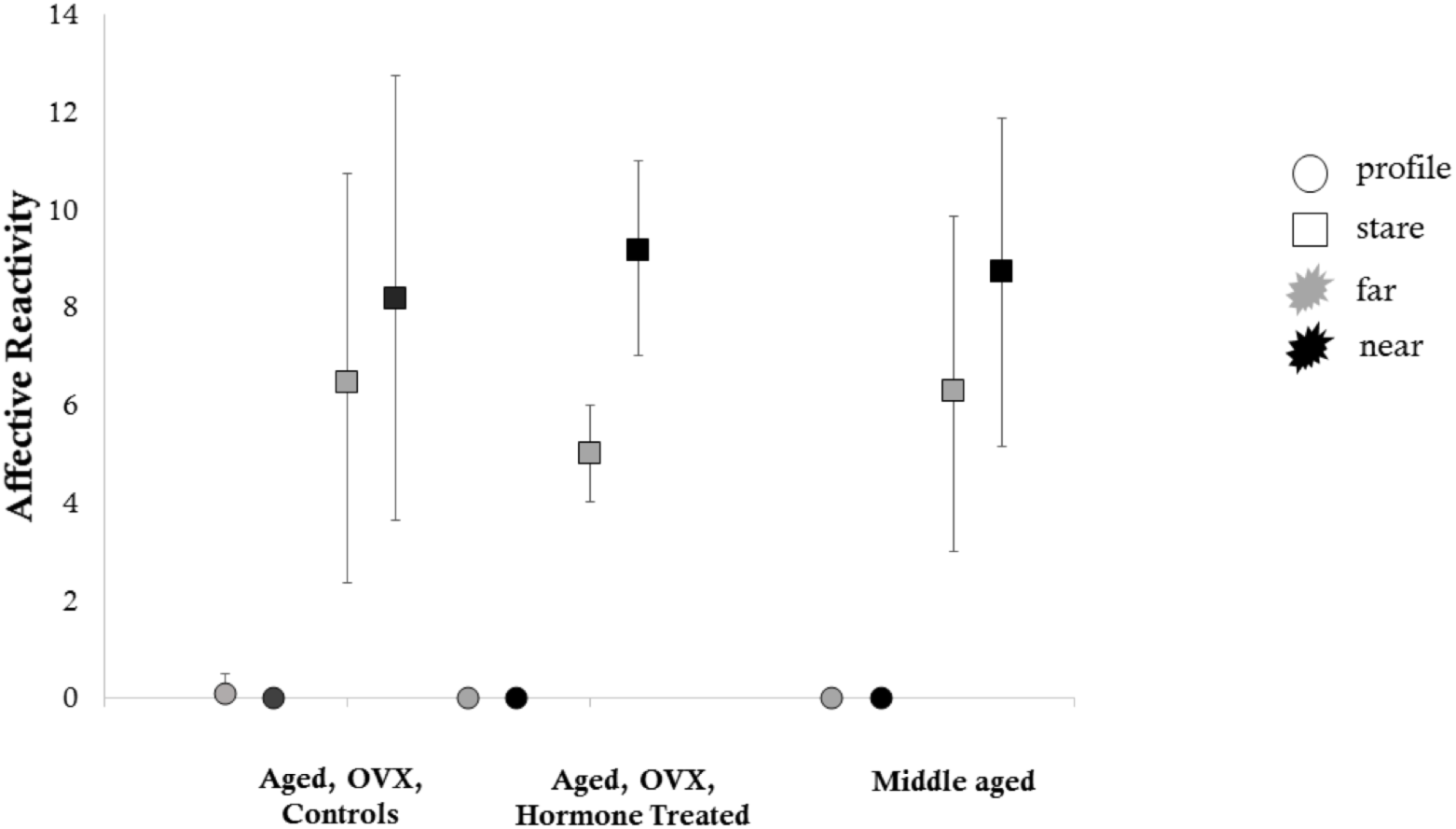
Mean affective reactitivy during the Human Intruder Task. Error bars represent bias corrected bootstrapped 95% confidence intervals. Circles indicate the profile condition. Squares indicate the stare condition. Grey filled shapes indicate the far condition. Black filled shapes indicate the near condition.

Visual inspection of the data suggested that the while, as a group, middle aged animals’ affective reactivity did not differ significantly between the two stare conditions, insofar as the 95% confidence intervals overlapped, individual middle aged animals appeared to be calibrating their responses and were more reactive during *stare-near* as compared to *stare-far*. We therefore computed a difference score for each animal between *stare-near* and *stare-far* for each condition and then computed bias corrected boot strapped means and 95% confidence intervals around those means. The 95% confidence interval around the mean difference score for the aged control animals included 0, indicating that these animals were not more reactive during *stare-near* than *stare-far*. In contrast, the 95% confidence interval around the mean difference score for the aged hormone treated animals and the middle aged animals did not include 0, indicating that these animals were significantly more reactive during *stare-near* than *stare-far*. That is, hormone treatment preserved middle aged affective reactivity. See Figure 2.

**Figure 2:**
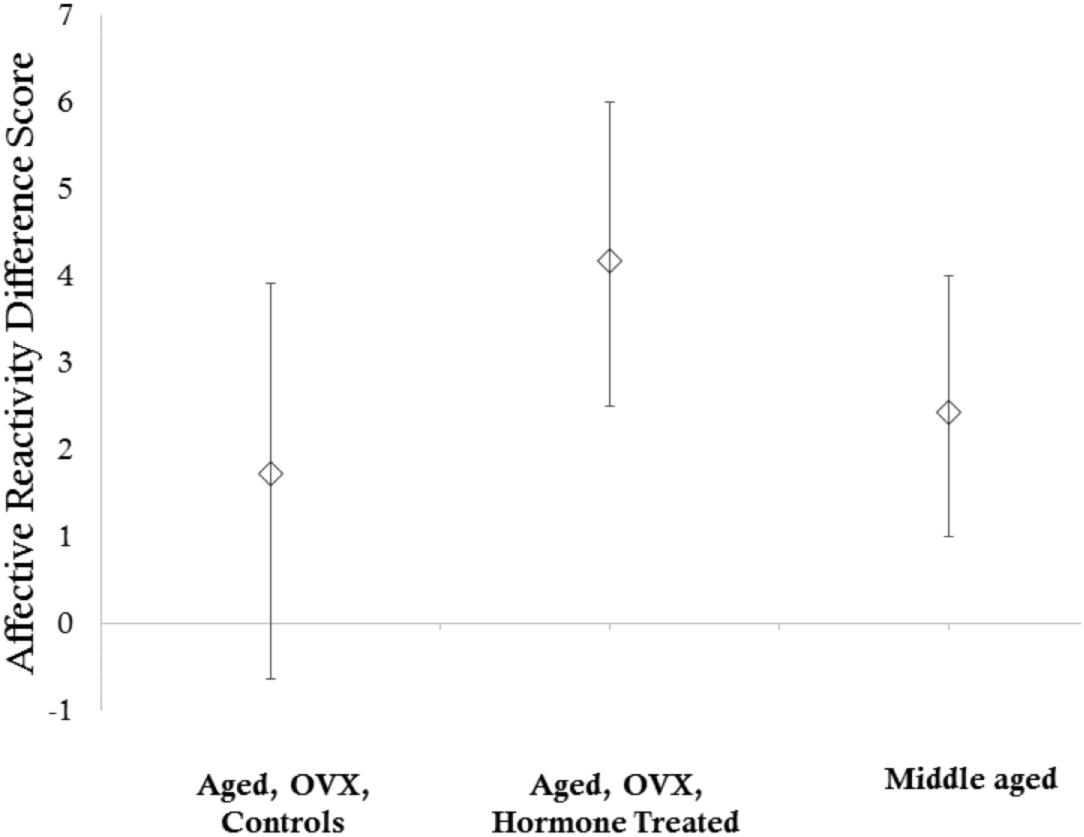
Mean affective reactitivy difference scores during the Human Intruder Task. Difference scores were computed by subtracting affective reactivity during the stare-far condition from affective reactivity during the stare-near condition for each individual subject. Error bars represent bias corrected bootstrapped 95% confidence intervals. Note that the overlapping confidence intervals of the three groups indicate that average affective reactivity was consistent across groups.

## Discussion

The present study demonstrates that responsivity to threat changes with age in rhesus monkeys, but not in the way predicted by the positivity effect. The positivity effect (Carstensen, Isaacowitz, & Charles, 1999; Carstensen, 2006; Mather & Carstensen, 2005; Reed & Carstensen, 2012) effect in humans demonstrates that as humans age emotional life becomes less negative and more positive. This would predict that animals serving as a model normal human female aging absent medical intervention – aged rhesus monkeys who had undergone surgical menopause and were not on hormone replacement therapy – would be less responsive to a potent threat of an unfamiliar human making eye contact. Instead, levels of affective reactivity were comparable in aged and middle aged monkeys. What did emerge, however, was that compared to middle aged monkeys, aged OVX non-hormone treated monkeys did not calibrate their behavioral responses to the magnitude of threat presented to them. Instead, they were statistically equally responsive to both moderate and potent threats. Hormone treatment with estradiol eliminated this effect. Aged OVX hormone-treated animals, like middle aged animals, calibrated their responses to the magnitude of threat, showing greater affective reactivity to the potent as compared to moderate threat. Additionally, aged OVX animals on hormone treatment as a group evidenced less variance in their responses, compared to both aged OVX animals not on hormone treatment and middle aged animals. These findings suggest that hormone treatment post-menopause may preserve some aspects of affective functioning typical of younger individuals.

What is not clear from our study is the extent to which variation in circulating levels of hormones impacted individual monkeys' affective reactivity. Some evidence from the human literature (for reviews Gasbarri et al., 2012; Georgakis et al., 2016; Newhouse & Albert, 2015; Sherwin, 1996) and minimal evidence from the nonhuman primate literature (e.g., Lacreuse & Herndon, 2003; Lacreuse et al., 2007) suggests that the levels of circulating estrogen may have an impact on affective behavior, although the specific patterns of effects are not entirely clear. Evaluating individual differences in hormone levels as they relate to individual differences in affective reactivity, both in aged OVX animals on hormone treatment and in middle aged animals with normal estrogen production, is a clear future direction for this program of work.

Our results demonstrate that aged, surgically menopausal female monkeys have altered patterns of affective behavior compared to middle-aged monkeys, insofar as they do not appropriately calibrate their affective responses. Middle-aged monkeys and aged surgically menopausal monkeys on estradiol treatment showed the same patterns that have been widely documented: as threat levels increase, so too does affective responding. Despite wide interest in human healthy affective aging, animal studies have primarily paid attention to cognition (for reviews Baxter, 2001; Voytko & Tinkler, 2004) and disease models (for reviews (Colman, 2017; Havel, Kievit, Comuzzie, & Bremer, 2017). Only very recently have nonhuman primates’ affective processes been considered in the context of aging (Almeling, Hammerschmidt, Sennhenn-Reulen, Freund, & Fischer, 2016; Weiss, King, Inuoue-Murayama, Matsuzawa, & Oswald, 2012). Understanding the normal, healthy, affective lives of nonhuman primates as they age is critical for establishing them as a model for human age-related processes in which affect and emotion play a key role. While most humans’ affective lives fare well with age, some do not – elder depression and social isolation can have serious consequences (Alexopoulos, 2005; Beekman, Deeg, Braam, Smit, & Van Tillburg, 1997; Cornwell & Waite, 2009; Gerst-Emerson & Jayawardhana, 2015; Valtorta & Hanratty, 2012). Further, many age-related diseases such as Alzheimer’s disease have comorbid mood and affect problems, the mechanisms of which are not well characterized. The assessment of affective reactivity in aged monkeys, and the documentation of a marked effect of hormone treatment in aged surgically menopausal monkeys, is a first step towards the incorporation of affect and emotion in experimental studies in this species. We anticipate that this model will provide new insights into the neurobiological and hormonal mechanisms of affective aging in a species that has been a particularly effective model of human cognitive and reproductive aging (Hara et al., 2014; Morrison & Baxter, 2012).

## Authors’ Note

Eliza Bliss-Moreau, Department of Psychology, University of California, Davis, and California National Primate Research Center; Mark G. Baxter, Department of Neuroscience and Friedman Brain Institute, Icahn School of Medicine at Mount Sinai.

This project was supported by National Institute on Aging (NIA) award P01-AG016765. EBM was supported by K99-MH10138 during data collection. The California National Primate Research Center is supported by National Institutes of Health (NIH) Office of the Director award P51-OD011107. The content is solely the responsibility of the authors and does not necessarily represent the official views of the NIH.

We thank Gilda Moadab, Mary Roberts, Tracy Ojakangas, and Lisa Novik for technical assistance, and Nancy Gee and Bill Lasley for hormone assay data verifying ovariectomy.

## References

Alexopoulos, G. S. (2005). Depression in the elderly. The Lancet, 365(9475), 1961–1970. https://doi.org/10.1016/S0140-6736(05)66665-2

Almeling, L., Hammerschmidt, K., Sennhenn-Reulen, H., Freund, A. M., & Fischer, J. (2016). Motivational Shifts in Aging Monkeys and the Origins of Social Selectivity. Curr Biol, 26(13), 1744–1749. https://doi.org/10.1016/j.cub.2016.04.066

Baxter, M. G. (2001). Cognitive aging in nonhuman primates. In P. R. Hoff & C. V Mobbs (Eds.), Functional Neurobiology of Aging (pp. 407–419). San Diego, CA: Academic Press. https://doi.org/10.1016/B978-012351830-9/50028-7

Baxter, M. G., Roberts, M. T., Gee, N. A., Lasley, B. L., Morrison, J. H., & Rapp, P. R. Multiple clinically relevant hormone therapy regimens fail to improve cognitive function in aged ovariectomized rhesus monkeys, 34 Neurobiol Aging § (2013). Elsevier Ltd. Retrieved from http://dx.doi.org/10.1016/j.neurobiolaging.2012.12.017

Baxter, M. G., Roberts, M. T., Gee, N. A., Lasley, B. L., Morrison, J. H., & Rapp, P. R. (2013b). Multiple clinically relevant hormone therapy regimens fail to improve cognitive function in aged ovariectomized rhesus monkeys. Neurobiol Aging, 34(7), 1882–1890. https://doi.org/10.1016/j.neurobiolaging.2012.12.017

Beekman, A., Deeg, D., Braam, A., Smit, J., & Van Tillburg, W. (1997). Consequences of major and minor depression in later life: A study of disability, well-being and service utilization. Psychological Medicine, 27(6), 1397–1409.

Bliss-Moreau, E., & Moadab, G. (2016). Variation in Behavioral Reactivity Is Associated with Cooperative Restraint Training Efficiency. J Am Assoc Lab Anim Sci, 55(1), 41–49. Retrieved from https://www.ncbi.nlm.nih.gov/pmc/articles/PMC4747010/pdf/jaalas2016000041.pdf

Carlsson, H. E., Schapiro, S. J., Farah, I., & Hau, J. (2004). Use of primates in research: a global overview. Am J Primatol, 63(4), 225–237. https://doi.org/10.1002/ajp.20054

Carstensen, L. L. (2006). The influence of a sense of time on human development. Science (New York, N.Y.), 312(5782), 1913–5. https://doi.org/10.1126/science.1127488

Carstensen, L. L., Isaacowitz, D. M., & Charles, S. T. (1999). Taking time seriously: A theory of socioemotional selectivity. American Psychologist, 54(3), 165–181.

Carstensen, L. L., & Mikels, J. A. (2005). At the intersection of emotion and cogntion. Current Directions in Psychological Science, 14(3), 117–121.

Carstensen, L. L., Pasupathi, M., Mayr, U., & Nesselroade, J. R. (2000). Emotional Experience in Everyday Life Across the Adult Life Span. Journal of Personality and Social Psychology, 79(4), 644–655.

Carstensen, L. L., Turan, B., Scheibe, S., Ram, N., Ersner-Hershfield, H., Samanez-Larkin, G. R.,…Nesselroade, J. R. (2011). Emotional experience improves with age: evidence based on over 10 years of experience sampling. Psychol Aging, 26(1), 21–33. https://doi.org/10.1037/a0021285

Chatterji, S., Byles, J., Cutler, D., Seeman, T., & Verdes, E. (2015). Health, functioning, and disability in older adults—present status and future implications. The Lancet, 385(9967), 563–575. https://doi.org/10.1016/s0140-6736(14)61462-8

Colman, R. J. (2017). Non-human primates as a model for aging. Biochimica et Biophysica Acta (BBA) – Molecular Basis of Disease. https://doi.org/10.1016/J.BBADIS.2017.07.008

Cornwell, B., Laumann, E. O., & Schumm, L. P. (2008). The Social Connectedness of Older Adults: A National Profile. Am Sociol Rev, 73(2), 185–203.

Cornwell, E. Y., & Waite, L. J. (2009). Social Disconnectedness, Perceived Isolation, and Health among Older Adults. Journal of Health and Social Behavior, 50(1), 31–48. https://doi.org/10.1177/002214650905000103

Frewen, P. A., Dozois, D. J., Joanisse, M. F., & Neufeld, R. W. (2008). Selective attention to threat versus reward: meta-analysis and neural-network modeling of the dot-probe task. Clin Psychol Rev, 28(2), 307–337. https://doi.org/10.1016/j.cpr.2007.05.006

Gasbarri, A., Tavares, M. C., Rodrigues, R. C., Tomaz, C., & Pompili, A. (2012). Estrogen, cognitive functions and emotion: an overview on humans, non-human primates and rodents in reproductive years. Rev Neurosci, 23(5-6), 587–606. https://doi.org/10.1515/revneuro-2012-0051

Georgakis, M. K., Thomopoulos, T. P., Diamantaras, A. A., Kalogirou, E. I., Skalkidou, A., Daskalopoulou, S. S., & Petridou, E. T. (2016). Association of Age at Menopause and Duration of Reproductive Period With Depression After Menopause: A Systematic Review and Meta-analysis. JAMA Psychiatry, 1–12. https://doi.org/10.1001/jamapsychiatry.2015.2653

Gerst-Emerson, K., & Jayawardhana, J. (2015). Loneliness as a public health issue: the impact of loneliness on health care utilization among older adults. American Journal of Public Health, 105(5), 1013–9. https://doi.org/10.2105/AJPH.2014.302427

Gottlieb, D. H., & Capitanio, J. P. (2013). Latent variables affecting behavioral response to the human intruder test in infant rhesus macaques (Macaca mulatta). Am J Primatol, 75(4), 314–323. https://doi.org/10.1002/ajp.22107

Guapo, V. G., Graeff, F. G., Zani, A. C., Labate, C. M., dos Reis, R. M., & Del-Ben, C. M. (2009). Effects of sex hormonal levels and phases of the menstrual cycle in the processing of emotional faces. Psychoneuroendocrinology, 34(7), 1087–1094. https://doi.org/10.1016/j.psyneuen.2009.02.007

Hara, Y., Yuk, F., Puri, R., Janssen, W. G. M., Rapp, P. R., & Morrison, J. H. Presynaptic mitochondrial morphology in monkey prefrontal cortex correlates with working memory and is improved with estrogen treatment., 111 Proceedings of the National Academy of Sciences § (2014). Retrieved from http://eutils.ncbi.nlm.nih.gov/entrez/eutils/elink.fcgi?dbfrom=pubmed&id=24297907&retmode=ref&cmd=prlinks

Havel, P. J., Kievit, P., Comuzzie, A. G., & Bremer, A. A. (2017). Use and Importance of Nonhuman Primates in Metabolic Disease Research: Current State of the Field. ILAR Journal, 58(2), 251–268. https://doi.org/10.1093/ilar/ilx031

Isaacowitz, D. M., Wadlinger, H. A., Goren, D., & Wilson, H. R. (2006a). Is there an age-related positivity effect in visual attention? A comparison of two methodologies. Emotion, 6(3), 511–516. https://doi.org/10.1037/1528-3542.6.3.511

Isaacowitz, D. M., Wadlinger, H. A., Goren, D., & Wilson, H. R. (2006b). Selective preference in visual fixation away from negative images in old age? An eye-tracking study. Psychol Aging, 21(1), 40–48. https://doi.org/10.1037/0882-7974.21.1.40

Kalin, N. H., & Shelton, S. E. Defensive behaviors in infant rhesus monkeys: environmental cues and neurochemical regulation., 243 Science § (1989). Retrieved from http://eutils.ncbi.nlm.nih.gov/entrez/eutils/elink.fcgi?dbfrom=pubmed&id=2564702&retmode=ref&cmd=prlinks

Kalin, N. H., & Shelton, S. E. (1998). Ontogeny and stability of separation and threat-induced defensive behaviors in rhesus monkeys during the first year of life. Am J Primatol, 44(2), 125–135. https://doi.org/10.1002/(sici)1098-2345(1998)44:2<125::aid-ajp3>3.0.co;2-y

Kamboj, S. K., Krol, K. M., & Curran, H. V. (2015). A specific association between facial disgust recognition and estradiol levels in naturally cycling women. PLoS One, 10(4), e0122311. https://doi.org/10.1371/journal.pone.0122311

Lacreuse, A., & Herndon, J. G. (2003). Estradiol selectively affects processing of conspecifics’ faces in female rhesus monkeys. Psychoneuroendocrinology, 28(7), 885–905. https://doi.org/10.1016/S0306-4530(02)00104-X

Lacreuse, A., Martin-Malivel, J., Lange, H. S., & Herndon, J. G. (2007). Effects of the menstrual cycle on looking preferences for faces in female rhesus monkeys. Animal Cognition, 10(2), 105–115. https://doi.org/10.1007/s10071-006-0041-8

Mather, M., & Carstensen, L. L. (2003). Aging and Attentional Biases for Emotional Faces. Psychological Science, 14(5), 409–415.

Mather, M., & Carstensen, L. L. (2005). Aging and motivated cognition: the positivity effect in attention and memory. Trends Cogn Sci, 9(10), 496–502. https://doi.org/10.1016/j.tics.2005.08.005

McPherson, M., Smith-Lovin, L., & Brashears, M. E. (2006). Social Isolation in America: Chagnes in Core Discussion Networks over Two Decades. American Sociological Review, 71(3), 353–375.

Morrison, J. H., & Baxter, M. G. The ageing cortical synapse: hallmarks and implications for cognitive decline., 13 Nat Rev Neurosci § (2012). Retrieved from http://eutils.ncbi.nlm.nih.gov/entrez/eutils/elink.fcgi?dbfrom=pubmed&id=22395804&retmode=ref&cmd=prlinks

Newhouse, P., & Albert, K. (2015). Estrogen, Stress, and Depression: A Neurocognitive Model. JAMA Psychiatry, 72(7), 727–729. https://doi.org/10.1001/jamapsychiatry.2015.0487

Nikitin, J., & Freund, A. M. (2011). Age and motivation predict gaze behavior for facial expressions. Psychol Aging, 26(3), 695–700. https://doi.org/10.1037/a0023281

Rapp, P. R., Morrison, J. H., & Roberts, J. A. Cyclic estrogen replacement improves cognitive function in aged ovariectomized rhesus monkeys, 23 J Neurosci § (2003). Retrieved from http://www.jneurosci.org/content/23/13/5708.full.pdf

Raz, N. (2000). Aging of the brain and its impact on cognitive performance: Integration of structural and functional findings. In F. I. M. Craik & T. A. Salthouse (Eds.), Handbook of Aging and Cognition – II (pp. 1–90). Mahwah, NJ: Erlbaum.

Reed, A. E., & Carstensen, L. L. (2012). The theory behind the age-related positivity effect. Front Psychol, 3, 339. https://doi.org/10.3389/fpsyg.2012.00339

Sherwin, B. B. (1996). Hormones, mood, and cognitive functioning in postmenopausal women. Obstet Gynecol, 87(2 Suppl), 20s–26s.

Sherwin, B. B., & Disord, J. A. (1988). Affective changes with estrogen and androgen replacement therapy in surgically menopausal women. Journal of Affective Disorders, 14(2), 177–187.

Valtorta, N., & Hanratty, B. (2012). Loneliness, isolation and the health of older adults: do we need a new research agenda? Journal of the Royal Society of Medicine, 105(12), 518–522. https://doi.org/10.1258/jrsm.2012.120128

Voytko, M. L., & Tinkler, G. P. (2004). Cognitive function and its neural mechanisms in nonhuman primate models of aging, Alzheimer disease, and menopause. Frontiers in Bioscience: A Journal and Virtual Library, 9, 1899–914. Retrieved from http://www.ncbi.nlm.nih.gov/pubmed/14977596

Weiss, A., King, J. E., Inuoue-Murayama, M., Matsuzawa, T., & Oswald, A. J. (2012). Evidence for a midlife crisis in great apes consistent with the U-shape in human wellbeing. Proceedings of the National Academy of Science, 109(49), 19949–19952.

